# Transcriptional Blood Signatures for Active and Amphotericin B Treated Visceral Leishmaniasis in India

**DOI:** 10.1101/554022

**Authors:** Michaela Fakiola, Om-Prakash Singh, Genevieve Syn, Toolika Singh, Bhawana Singh, Jaya Chakravarty, Shyam Sundar, Jenefer M. Blackwell

## Abstract

Amphotericin B provides improved therapy for visceral leishmaniasis (VL) caused by *Leishmania donovani*, with single dose liposomal-encapsulated Ambisome providing the best cure rates. The VL elimination program aims to reduce the incidence rate in the Indian subcontinent to <1/10,000 population/year. Ability to predict which asymptomatic individuals (e.g. anti-leishmanial IgG and/or Leishmania-specific modified Quantiferon positive) will progress to clinical VL would help in monitoring disease outbreaks. Here we examined whole blood transcriptional profiles associated with asymptomatic infection, active disease, and in treated cases. Two independent microarray experiments were performed, with analysis focussed primarily on differentially expressed genes (DEGs) concordant across both experiments. No DEGs were identified for IgG or Quantiferon positive asymptomatic groups compared to negative healthy endemic controls. We therefore concentrated on comparing concordant DEGs from active cases with all healthy controls, and in examining differences in the transcriptome following different regimens of drug treatment. In these comparisons 6 major themes emerged: (i) expression of genes and enrichment of gene sets associated with erythrocyte function in active cases; (ii) strong evidence for enrichment of gene sets involved in cell cycle in comparing active cases with healthy controls; (iii) identification of *IFNG* encoding interferon-γ as the major hub gene in concordant gene expression patterns across experiments comparing active cases with healthy controls or with treated cases; (iv) enrichment for interleukin signalling (IL-1/3/4/6/7/8) and a prominent role for CXCL10/9/11 and chemokine signalling pathways in comparing active cases with treated cases; (v) the novel identification of Aryl Hydrocarbon Receptor signalling as a significant canonical pathway when comparing active cases with healthy controls or with treated cases; and (vi) global expression profiling support for more effective cure at day 30 post-treatment with a single dose of liposomal encapsulated amphotericin B compared to multi-dose non-liposomal amphotericin B treatment over 30 days.

**Author Summary:** Visceral leishmaniasis (VL), also known as kala-azar, is a potentially fatal disease caused by intracellular parasites of the *Leishmania donovani* species complex. VL is a serious public health problem in rural India, causing high morbidity and mortality, as well as major costs to local and national health budgets. Amphotericin B provides improved therapy for VL with single dose liposomal-encapsulated Ambisome, now affordable through WHO-negotiated price reductions, providing the best cure rates. The VL elimination program aims to reduce the incidence rate in the Indian subcontinent to <1/10,000 population/year. By assessing immune responses to parasites in people infected with *L. donovani*, but with different clinical status, we can determine the requirements for immune cell development and predict which asymptomatic individuals, for example healthy individuals with high anti-leishmanial antibody levels, will progress to clinical VL. This will help in monitoring disease outbreaks. In this study we looked at global gene expression patterns in whole blood to try to understand more about asymptomatic infection, active VL, and the progress to cure in cases treated with single or multi-dose amphotericin B. The signatures of gene expression identified aid in our understanding of disease pathogenesis and provide novel targets for therapeutic intervention in the future.

## Introduction

Visceral leishmaniasis (VL), also known as kala-azar, is a potentially fatal disease caused by obligate intracellular parasites of the *Leishmania donovani* species complex. VL is a serious public health problem in indigenous and rural populations in India, causing high morbidity and mortality, as well as major costs to both local and national health budgets. The estimated annual global incidence of VL is 200,000 to 400,000, with up to 50,000 deaths annually occurring principally in India, Bangladesh, Sudan, South Sudan, Ethiopia and Brazil [1]. In India, improvements in drug therapy have been afforded through the introduction of amphotericin B treatment, with single dose liposomal encapsulated Ambisome providing the best cure rates and now being used as the preferred treatment regime in the VL elimination program [2]. However, with the potential development of drug resistance to each new therapeutic approach [3], there remains a continuing need for improved and more accurate methods of early diagnosis, as well as ability to monitor responses to treatment and to predict disease outcome. These objectives are also important in relation to the World Health Organization-supported VL elimination initiative in the Indian subcontinent, which aims at reducing the incidence rate of VL in the region to below 1 per 10,000 population per year by 2020 [4]. Monitoring disease outbreaks in the context of the elimination program will be an important goal, including the ability to determine which individuals displaying asymptomatic disease, as monitored by anti-leishmanial IgG [5,6] and/or Leishmania-specific modified Quantiferon responses [7], will progress to clinical VL disease [6].

In recent years, the use of whole blood transcriptional profiling has provided a better understanding of the host response to infectious disease, leading to the identification of blood signatures and potential biomarkers for use in diagnosis, prognosis and treatment monitoring (reviewed [8]). This approach was successful in identifying a neutrophil-driven interferon (IFN)-inducible blood transcriptional signature for active tuberculosis that involved both IFN-γ and type I IFN-α/β signalling [9] and was subsequently confirmed in multiple countries world-wide (reviewed [8]). This neutrophil-driven interferon signature was present in active disease but absent in both latent infection and in healthy controls [9]. While an IFN-inducible signature was also identified in patients with the autoimmune disease systemic lupus erythematosus, there were differences in the signatures that also distinguished the two profiles from each other [9]. Viral infections [10] and bacterial infections like melioidosis [11] are also broadly characterised by IFN-inducible gene expression, but whole blood signatures have been identified that are able to discriminate between bacterial and viral infections [10,12], as well as between different viral infections [10]. In HIV, blood transcriptional signatures have been identified that distinguish between rapid compared to slow progression to disease [13]. Blood signatures have also been identified which distinguish between children who acquire dengue virus fever compared to those who develop dengue haemorrhagic fever [14,15]. There are also signatures that distinguish between pulmonary and extra-pulmonary tuberculosis [16], as well as between pulmonary tuberculosis, pulmonary sarcoidosis, pneumonias and lung cancers [17]. A transcriptional signature that can be used to monitor treatment response is also a valuable goal in infectious disease. Again, studies from two cohorts followed longitudinally in South Africa show that the transcriptional signature of active tuberculosis disease rapidly diminishes with successful treatment [18,19].

In this study we set out to determine whole blood transcriptional profiles that might distinguish asymptomatic infection with *L. donovani* from active disease, as well as to monitor the changes in transcriptional profiles that accompanied drug treatment. Whilst we were unable to detect a signature that distinguished asymptomatic (IgG antibody positive [5,6], or modified Quantiferon positive [7]) individuals from healthy endemic controls who were negative by these two assays, we were able to determine the transcriptional profile of active VL cases, and to demonstrate interesting differences in return to baseline between patients treated with non-liposomal compared to liposomal-encapsulated (Ambisome™) amphotericin B.

## Methods

### Study Subjects

In this study two independent microarray experiments were performed. For experiment 1, samples were collected between February and April 2011. For experiment 2, samples were collected between April and July 2012. Samples were collected at the Kala-azar Medical Research Center (KAMRC), Muzaffarpur, Bihar, India, or in nearby field sites for some asymptomatic individuals and endemic controls. Active VL cases were diagnosed by experienced clinicians based on clinical signs, including fever (>2 weeks), splenomegaly, positive serology for recombinant antigen (r)-K39 and/or by microscopic demonstration of *Leishmania* amastigotes in splenic aspirate smears. VL patients were treated according to routine clinical care with either (a) experiment 1: 0.75 mg/kg non-liposomal amphotericin B daily for 15 days (N=3), or on alternate days over 30 days (N=7), by infusion (i.e. 15 doses in all; total dose 11.25 mg/kg over 30 days); or (b) experiment 2: 10 mg/kg of Ambisome (liposome-encapsulated amphotericin B) as a single dose by infusion. Blood samples were collected pre-(N=10 experiment 1; N=11 experiment 2) and post-(day 30; N=10 experiment 1; N=11 experiment 2) treatment. There were 9 paired pre-/post-treatment samples for experiment 1; 10 for experiment 2. Healthy control subjects included (i) asymptomatic individuals (N=2 experiment 1; N=6 experiment 2) who had high anti-leishmanial antibody levels by direct agglutination test (DAT titer ≥1:25,600) [6]; (ii) asymptomatic individuals who were positive by *Leishmania*-specific modified quantiferon assays [7] (N=8 experiment 1; N=9 experiment 2); and (iii) Serology (DAT titer ≤1:1600) and quantiferon negative healthy endemic controls (N=6 experiment 1; N=10 experiment 2). Sample sizes are for post-QC samples used in expression profiling studies (cf. below). Further clinical and demographic details on participants are provided in S1 Table.

### Ethics

The enrolment of human subjects complies with the principles laid down in the Helsinki declaration. Institutional ethical approval (reference numbers: Dean/2012-2013/89) was obtained from the ethical review board of Banaras Hindu University (BHU), Varanasi, India. Informed written consent was obtained from each participant at the time of enrolment, or from their legal guardian if they were under 18 years old. Only patients who had not previously received treatment and who agreed to participate in the study were enrolled. All clinical treatment and follow-up records were maintained using standardised case report forms on an electronic server. All patient data were analysed anonymously.

### RNA extraction and microarray analysis

Whole blood (5 mL) collected by venepuncture was immediately placed into Paxgene tubes (QIAGEN GmbH, Germany) and stored at −80°C for later processing for RNA. RNA was extracted using PAXgene Blood RNA kits (QIAGEN GmbH, Germany) according to manufacturer’s instructions. RNA integrity and purity were checked using Tape Station 4200 (Agilent Technologies, USA). Only RNAs with RNA integrity (RIN) values >5.5 were taken forward for beadchip analysis. Globin mRNA was depleted using GLOBINclear™-Human kits (ThermoFischer Scientific, USA). RNA was reverse transcribed and biotin-labelled using the Illumina TotalPrep™ RNA Amplification kit (ThermoFischer Scientific, USA). The resulting biotinylated cRNA was hybridised to Illumina HT12v4 Expression BeadChips, specifically HumanHT-12_V4_0_R2_15002873_B, containing 47,323 genome wide gene probes, and 887 control probes. Samples from different control or clinical groups were distributed evenly across 3 (experiment 1) or 4 (experiment 2) beadchips. All RNA preparation and processing of samples over beadchips was carried out at Sandor Lifesciences Pvt. Ltd. (Hyderabad, India).

### Data analysis

All data analysis was carried out in R Version 3.4.3 (Smooth Sidewalk - https://www.rproject.org/) and RStudio (version 1.1.383). The Bioconductor package *Lumi* [20] was used to read in raw expression values and perform quality control. Background correction and quantile normalisation of the data was carried out using the Bioconductor package *Limma* [21]. Pre-processing of the microarray data and removal of non-expressed (detection *P*-value > 0.05 in all arrays) and poor quality probes previously shown to have unreliable annotation [22] provided 21,959 and 23,466 probe sets which passed QC in experiments 1 and 2, respectively. Principal components analysis (PCA) and unsupervised cluster analysis (Pearson’s correlation coefficient; hclust=complete) of normalised data was performed in R. Data was visualised using the R packages *ggplot2* (3.1.2) [23] and *pheatmap* (1.0.12) [24]. Differential expression analysis using linear modelling and empirical Bayes methods was carried out in the Bioconductor package *Limma* [21] for comparisons between control and clinical groups, as indicated. The threshold for differential expression was a log_2_-fold-change ≥0 (i.e. ≥2-fold) and/or Benjamini-Hochberg [25] adjusted p-value (*P*_adj_) ≤0.05, as indicated. Genes achieving these thresholds were taken forward in analyses using the gene set enrichment tool Enrichr [26], and using Ingenuity Pathway Analysis (IPA) (Ingenuity^®^ Systems, www.ingenuity.com) to identify canonical pathways, upstream regulators, and gene networks. Enrichr [26,27] accesses a wide range of open access databases to identify pathways for which the gene set is enriched. It uses four scores to report enrichment: a p-value (reported here as *P*_nominal_) calculated using Fisher’s exact test; a q-value (reported here as *P*_adj_) which is the Benjamin-Hochberg adjusted p-value; a rank or z-score computed using a modification to Fisher’s exact test to compute deviation from an expected rank; and a combined score which is a combination of the p-value and z-score calculated by multiplying the two scores using the formula c=ln(p)*z. IPA uses the Ingenuity Knowledge Base, an extensive database comprising biological pathways and functional annotations derived from the interactions between genes, proteins, complexes, drugs, tissues and disease, to carry out all its analyses. Benjamini-Hochberg correction was applied where applicable and *P*_adj_ ≤ 0.05 was used to filter all results. Upstream Regulator Analysis within IPA was employed to predict if there were any endogenous genes/cytokines/transcription factors which may be responsible for the observed gene expression patterns. If an upstream regulator is identified, an activation Z-score is calculated based on the fold change values of its target genes within the dataset. A Z-score ≥2 suggests that an upstream regulator is activated whereas a Z-score ≤-2 suggests it is inhibited, with active VL cases being the experimental group of baseline comparator. IPA also generates a “Top Tox List” pathway which provides an indication of toxic or pathogenic pathways that could be amenable to therapeutic intervention. Networks were constructed in IPA using the “Connect” option under the “Build” functionality. Genes with no previously documented interactions were removed from the diagram and the functions of each network were inferred from the remaining connected genes in each time-point.

## Results

### Comparative transcriptomics across clinical groups

Two independent microarray experiments were carried out to compare transcriptional profiles across clinical groups that included active VL cases pre-treatment (N=10 experiment 1; N=11 experiment 2), drug treated VL cases (N=10 experiment 1; N=11 experiment 2), modified quantiferon [7] positive asymptomatic individuals (N=8 experiment 1; N=9 experiment 2), high Leishmania-specific antibody positive (by DAT) asymptomatic individuals (N=2 experiment 1; N=6 experiment 2), and endemic healthy controls (N=6 experiment 1; N=10 experiment 2) who were both modified quantiferon negative and antibody negative by DAT. PCA of the top 500 most variable probes (Fig 1) across all pairwise comparisons of samples showed that principal component 1 (PC1) accounted for 45% (experiment 1; Fig 1A and B) and 31% (experiment 2; Fig 1D and E) of the variation and resolved active cases compared to endemic healthy control and asymptomatic groups. The latter were not well resolved from each other in either experiment. Treated patients sat intermediate between, and overlapping with, both active cases and control/asymptomatic groups in experiment 1 but showed greater overlap with control/asymptomatic groups in experiment 2 (cf. below). This is particularly apparent when comparing plots of PC1 by PC3 (Fig 1B and 1E). Unsupervised hierarchical cluster analysis also (Fig 1C and 1F) provided discrete clusters of active cases compared to control and asymptomatic individuals, with treated cases interspersed with both active cases and control groups and not falling into a single discrete cluster in either experiment.

**Figure 1.**
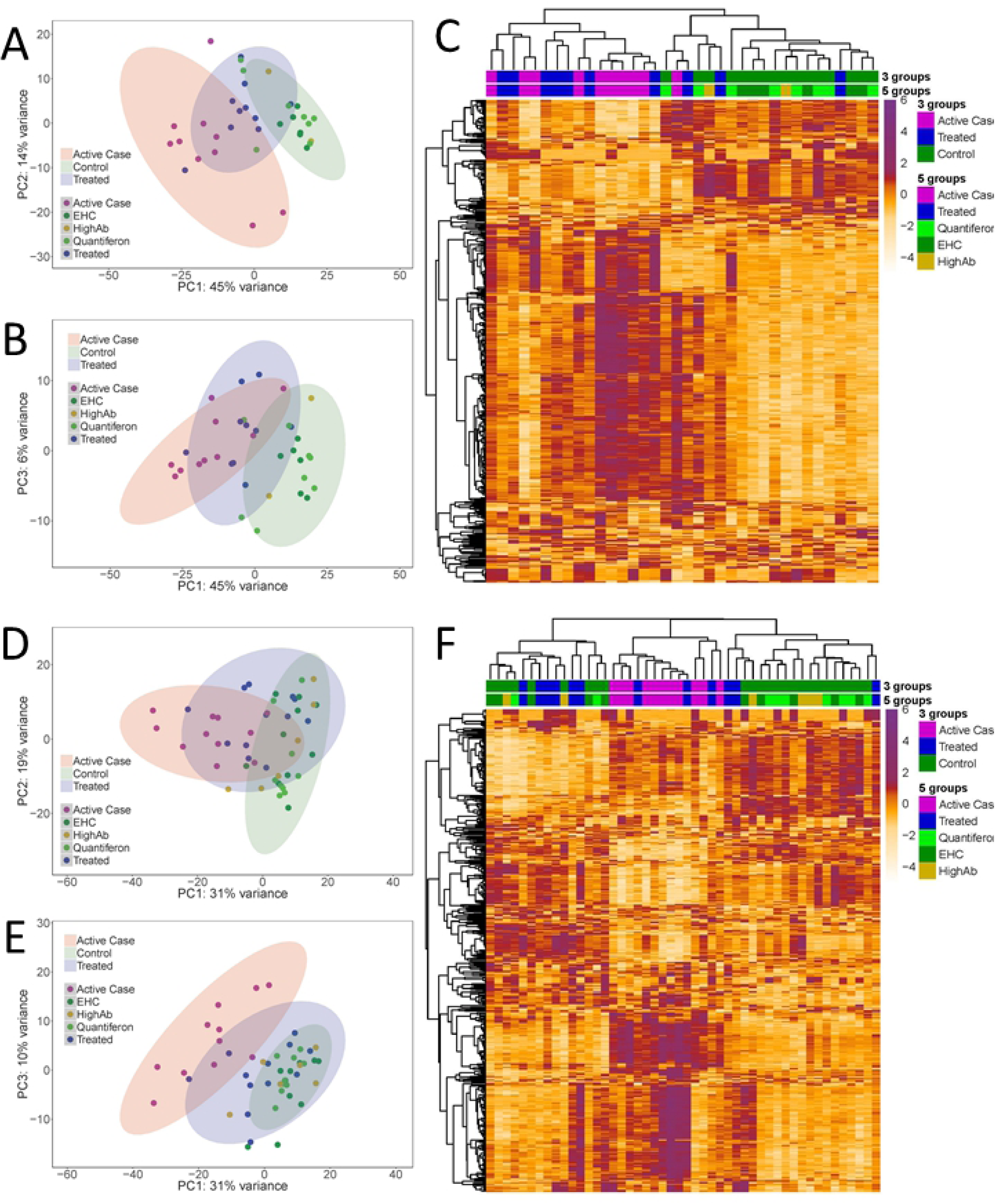
Principal Components Analysis (PCA) and hierarchical clustering of top 500 most variable probes. (A) PC1 by PC2 and (B) PC1 by PC3 in experiment 1, and (D) PC1 by PC2 and (E) PC1 by PC3 in experiment 2. Z-score transformed expression levels of the 500 most variable probes across all samples are represented as a heatmap for (C) experiment 1 and (F) experiment 2. Hierarchical clustering results based on Pearson’s correlation are shown as dendrograms on the top and left side of the matrix. Columns represent individual samples and rows individual probes. Experimental groups are color coded on the upper part of the heatmap. Active (=case) and treated (=treated) cases, as well as aymptomatics (=Quantiferon or HighAb positive individuals) and endemic healthy controls (=EHC), are colour coded as per the keys provided.

Consistent with the PCA plots (Fig 1), there were no differentially expressed probes representing genes (i.e. Benjamini-Hochberg [25] *P*_adj_ ≤0.05) when comparing either modified quantiferon positive asymptomatic individuals with endemic healthy controls, or when comparing high antibody titer individuals with endemic healthy controls, in either experiment 1 or experiment 2 (data not shown). These groups were therefore analysed as one group referred to as “controls” or “healthy controls” in all further differential expression analyses.

Differential expression analysis focused on the comparison of (i) active VL cases *versus* controls, (ii) treated VL cases *versus* controls, and (iii) active *versus* treated VL cases. Log_2_-fold-change in experiment 1 was highly correlated with log_2_-fold-change in experiment 2 across all probes (data not shown). Table 1 shows the number of differentially expressed probes representing genes in experiment 1 and experiment 2 for each comparison as well as the number of differentially expressed probes that replicated and were concordant for direction of effect between the two cohorts. At *P*_adj_ ≤0.05, there are 2,584 concordant differentially expressed probes in common when comparing active cases with controls, 37 concordant probes when comparing treated cases with controls, and 221 concordant probes when comparing active and treated cases. At the more stringent threshold of ≥2-fold change there were 439, 8, and 42 concordant probes for these comparisons, respectively.

**Table 1.** Summary of numbers of between group DEGs. DEGs at adjusted *P*-value ≤ 0.05 (top panel) and fold-change of expression ≥ 2 (bottom panel) for the comparison of the three main phenotype groups.

| **adj.$P \leq 0.05$** | **Experiment 1** | **Experiment 2** | **Concordant** |
| --- | --- | --- | --- |
| Case *vs* control | 4596 | 4651 | 2584 |
| Treated *vs* control | 1132 | 126 | 37 |
| Case *vs* treated | 654 | 1317 | 221 |

| **adj.$P \leq 0.05$**<br>**and fold-change $\geq 2$** | **Experiment 1** | **Experiment 2** | **Concordant** |
| --- | --- | --- | --- |
| Case *vs* control | 683 | 783 | 439 |
| Treated *vs* control | 120 | 27 | 8 |
| Case *vs* treated | 94 | 337 | 42 |

Of note, we found a greater number of transcriptional differences between treated cases and controls in experiment 1 compared to experiment 2 (Table 1; differentially expressed probes are 1132 and 126, respectively, at *P*_adj_ ≤0.05). One explanation for this could be the different treatment regimen employed in the two cohorts. VL patients of the first experiment were treated with 15 doses of a non-liposomal form of amphotericin B over 30 days. In experiment 2 patients received a single dose of liposomal amphotericin B, which has shown better efficacy for the treatment of VL [28,29]. The effect of treatment regimen on whole blood transcriptional profiles is further indicated by the comparison of active and treated cases. In this case, fewer differences in transcriptional regulation are observed between active and treated cases in experiment 1 as opposed to experiment 2 (Table 1; differentially expressed probes are 654 and 1317, respectively, at *P*_adj_ ≤0.05), in which patients have received a more efficacious therapy. These findings agree with the PCA results (Fig 1), in which treated cases of the experiment 1 cohort form a more discrete group between active cases and controls (Fig 1A and 1B) whereas treated cases of the experiment 2 cohort are grouped more closely to controls (Fig 1D and 1E).

Due to the small number of concordant differentially expressed genes identified for the treated cases *versus* controls (Table 1; 37 at *P*_adj_ ≤0.05, 8 at *P*_adj_ ≤0.05 and ≥2-fold change), only the concordant differentially expressed gene sets for active cases *versus* controls and active cases *versus* treated cases were used in subsequent pathway and gene set enrichment analyses.

### Network, pathway and gene set enrichment analyses comparing active cases and healthy controls

Heatmaps were generated for individual expression levels for probes representing the top 10 concordant genes expressed at a higher (“induced”) level (Fig 2A), and the top 10 concordant genes expressed at a lower (“repressed”) level (Fig 2B), in active cases compared to controls in experiment 1. Heatmaps for the same “induced” and “repressed” probes/genes in experiment 2 are presented in Fig 2C and 2D. Of note 8/10 “repressed” genes were also in the top 10 most highly differentially expressed “repressed” genes in experiment 2; all 10 genes achieved ≥2-fold change in both experiments. Amongst these 10 most “repressed” genes were: peptidase inhibitor 3 (*PI3*), a known antimicrobial peptide for bacteria and fungi that is upregulated by lipopolysaccharide and cytokines; the C-C chemokine ligand 23 (*CCL23*; represented by 2 probes) which acts as a chemoattractant for resting (but not active) T cells, monocytes, and to a lesser extent neutrophils; G-protein-coupled C-C motif chemokine receptor 3 (*CCR3*) which binds CCL10 (eotaxin), CCL26 (eotaxin-3), CCL7 (MCP3), CCL13 (MCP4) and CCL5 (RANTES) that likewise act as chemoattractants for eosinophils, monocytes and neutrophils; *ALOX15* which is a lipoxygenase known to regulate inflammation and immunity; and the G-protein-coupled prostaglandin D2 receptor 2 (*PTGDR2* alias *GPR44*) that is preferentially expressed in CD4 effector T helper 2 (Th2) cells and mediates pro-inflammatory chemotaxis of eosinophils, basophils and Th2 cells. For the “induced” genes (Fig 2A and 2C), only 2/10 (the top 2 in both experiments) were also in the top 10 “induced” genes in experiment 2, but all achieved ≥2-fold change in both experiments. In addition to type I interferon inducible 27 (*IFI27*) and complement C1q B chain (*C1QB*) genes, there was a bias amongst the most strongly “induced” genes towards genes involved in erythrocyte function, including: glycophorin B (*GYPB*), a major sialoglycoprotein of the human erythrocyte membrane; Rh D blood group antigens (*RHD*); hemoglobin subunit delta (*HBD*); 5’-aminolevulinate synthase 2 (*ALAS2*) an erythroid-specific enzyme located in the mitochondrion and involved in heme biosynthesis; carbonic anhydrase 1 (*CA1*) which is found at its highest level in erythrocytes; atypical chemokine receptor 1 (Duffy blood group) (*ACHR1* alias *DARC*) known for its role as the erythrocyte receptor for *Plasmodium vivax* and *P. knowlesi*; and 2,3-diphosphoglycerate (2,3-DPG) (*BPGM*) found at high concentrations in red blood cells where it binds to and decreases the oxygen affinity of haemoglobin.

**Figure 2.**
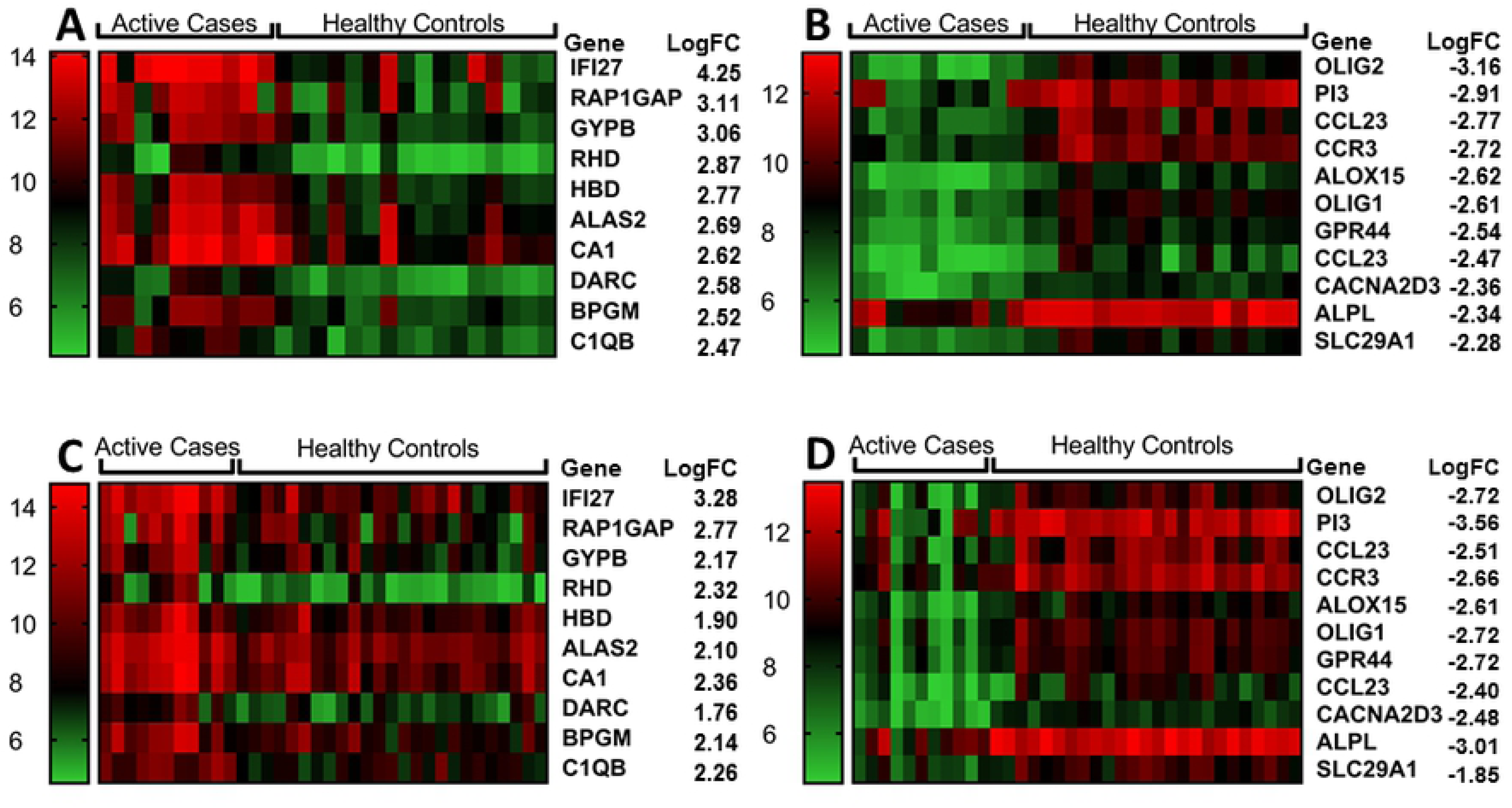
Heatmaps for top differentially expressed genes between active cases and healthy controls. (A) top 10 “induced” and (B) top 10 “repressed” genes for differential expression between active cases (N=10) and healthy controls (N=16) in experiment 1. (C) and (D) show heatmaps for the same genes in active cases (N=10) and healthy controls (N=25) using data from experiment 2. Columns represent individuals and rows represent individual genes, coloured to indicate expression levels based on post-QC normalised and log_2_-trasnformed data as indicated by the legend to the left of each figure. LogFC = log_2_ fold-change.

To gain a more global picture of the impact of differential gene expression, the 391 genes represented by 439 probes that were concordant for differential gene expression (*P*_adj_ ≤0.05; ≥2-fold change) between active cases and controls in experiments 1 and 2 were taken forward in Ingenuity Pathway (IPA) and gene-set enrichment (Enrichr) analyses. IPA network analysis indicated that 254 of these genes are joined in a single network (Fig 3), with *IFNG* as the major hub gene (i.e. with most connections to other genes in the network), and other major hub genes including *CCNA2, CXCL10, SPI1, SNCA, CHEK1, MCM2, AURKB, RARA, CDK1, CDC20,* and *FOXM1*. The top Ingenuity Canonical Pathways for the 391 genes that achieved *P*_adj_ <0.05 and ≥2-fold change (Table 2) were Estrogen-mediated S-phase Entry (*P*=6.46×10^-5^; *P*_adj_=0.019), Mitotic Roles of Polo-Like Kinases (*P*=1.20×10^-4^; *P*_adj_=0.019), Aryl Hydrocarbon Receptor (AHR) Signalling (*P*=1.51×10^-4^; *P*_adj_=0.019), and Heme Biosynthesis II (*P*=3.72×10^-4^; *P*_adj_=0.035). Although not achieving *P*_adj_≤0.05, identification of the Th2 pathway (*P*=1.22×10^-3^; *P*_adj_=0.074) and Activation of Th1 and Th2 Pathway (*P*=1.32×10^-3^; *P*_adj_=0.074) as nominally significant canonical pathways is consistent with prior knowledge of immune responses to leishmaniasis.

**Figure 3.**
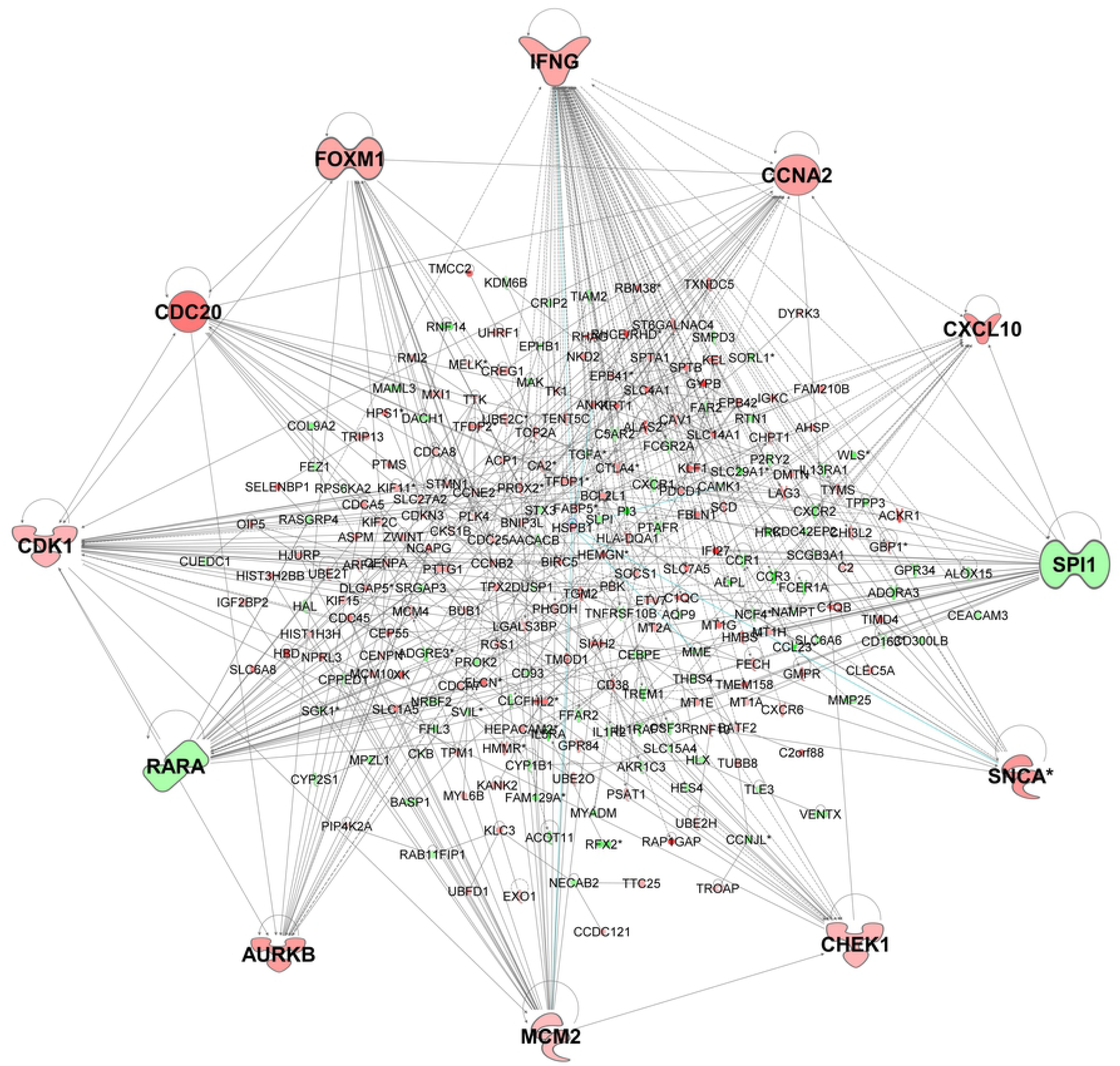
Gene network for concordant genes comparing active cases and healthy controls. The network was generated in IPA for 254 (of 391) genes concordant across experiments 1 and 2 for differential expression (adjusted p-value ≤0.05; ≥2-fold change) when comparing active cases and healthy controls. Genes in red have increased expression and genes in green have decreased expression when comparing active cases with healthy controls. The more intense the colour the larger the fold change values. Expression values are based in experiment 1, representative of similar results obtained for concordant genes across the two experiments.

**Table 2.** List of pathways identified by IPA Canonical Pathway Analysis. The table shows results for 439 probes representing 391 genes concordant for differential expression (adjusted P-value <0.05; >2-fold change) when comparing active VL cases with healthy controls across experiments 1 and 2, and for 221 probes representing 210 genes concordant for differential expression (adjusted P=value <0.05) when comparing active VL cases with treated VL cases across the two experiments.

| Ingenuity Canonical Pathways | $P_{\text{nominal}}$ | $P_{\text{adj}}$ | Genes |
| --- | --- | --- | --- |
| Active cases *versus* controls (391 top concordant genes; $P_{\text{adj}} < 0.05$; fold-change > 2) | | | |
| Estrogen-mediated S-phase Entry | $6.46 \times 10^{-5}$ | 0.019 | ↑CCNA2, ↑CCNE2, ↑TFDP1, ↑CDK1, ↑CDC25A |
| Mitotic Roles of Polo-Like Kinase | $1.20 \times 10^{-4}$ | 0.019 | ↑PLK4, ↑CDC20, ↑PTTG1, ↑CCNB2, ↑CDK1, ↑KIF11, ↑CDC25A |
| Aryl Hydrocarbon Receptor Signaling | $1.51 \times 10^{-4}$ | 0.019 | ↑TGM2, ↑CCNA2, ↑CCNE2, ↑NFIX, ↑TFDP1, ↓RARA, ↑ALDH5A1, ↓CYP1B1, ↑HSPB1, ↑CHEK1 |
| Heme Biosynthesis II | $3.72 \times 10^{-4}$ | 0.035 | ↑FECH, ↑ALAS2, ↑HMBS |
| Th2 Pathway | $1.12 \times 10^{-3}$ | 0.074* | ↓CCR1, ↑IFNG, ↓CCR3, ↓PTGDR2, ↑CXCR6, ↓PIK3R6, ↑HLA-DQA1, ↓SPI1, ↑TIMD4 |
| Th1 and Th2 Activation Pathway | $1.32 \times 10^{-3}$ | 0.074* | ↓CCR1, ↑IFNG, ↑SOCS1, ↓CCR3, ↓PTGDR2, ↑CXCR6, ↓PIK3R6, ↑HLA-DQA1, ↓SPI1, ↑TIMD4 |
| Active cases *versus* treated cases (210 Top concordant genes; $P_{\text{adj}} < 0.05$) | | | |
| Pathogenesis of Multiple Sclerosis | $9.33 \times 10^{-5}$ | 0.032 | ↑CXCL10, ↑CXCL11, ↑CXCL9 |
| Acute Myeloid Leukemia Signaling | $3.55 \times 10^{-4}$ | 0.049 | ↓CSF3R, ↑NRAS, ↓MAPK3, ↓RARA, ↓SPI1, ↓CSF1R |
| Oncostatin M Signaling | $4.37 \times 10^{-4}$ | 0.049 | ↑NRAS, ↓MAPK3, ↓CHI3L1, ↑STAT1 |
| Notch Signaling | $6.03 \times 10^{-4}$ | 0.05 | ↓MAML3, ↓LFNG, ↓RBPJ, ↓NOTCH1 |
| Thyroid Cancer Signaling | $7.41 \times 10^{-4}$ | 0.05 | ↑CXCL10, ↑NRAS, ↓MAPK3, ↓RXRA |
| Aryl Hydrocarbon Receptor Signaling | $3.09 \times 10^{-3}$ | 0.105** | ↑CCNE2, ↑NCOA7, ↓MAPK3, ↓RARA, ↓ALDH3B1, ↓RXRA |

\*Did not achieve adjusted P-value <0.05 but supported *a priori* by previous knowledge of immune response to leishmaniasis. \*\*Did not achieve adjusted P-value <0.05 but supports importance of the pathway identified by the comparison of active cases with healthy controls. Arrows indicated expression level in active cases relative to the relevant comparator.

Identification of AHR signalling as the top Ingenuity Canonical Pathway is reflective of increasing recognition of the role of AHR signalling in immunity, including the ability of AHR ligands to significantly induce cell secretion of IL-10 and inhibit IL-1β and IL-6 production in dendritic cells, and to promote IL-10 production and suppress IL-17 expression in CD4(+) T cells [30-32], and with the identification of *RARA, CCNA2* and *CHEK1* genes from the AHR pathway (Table 2) as major hub genes (Fig 3). AHR signalling was also identified as top in the Ingenuity “Top Tox List” pathway (*P*=4.16×10^-4^) indicative of its role as a toxic pathology endpoint that could be amenable to therapeutic intervention. Schematic representation of the core AHR canonical pathway overlaid with concordant gene expression data (*P*_adj_<0.05) for experiment 1 (Fig 4) for active cases relative to healthy controls shows differential gene expression that includes core players AHR and the AHR nuclear translocator (ARNT) in the AHR pathway, as well as for key phase I metabolising enzymes (CYPB1, ALDH5A1, ALD3B1 and ALD3A2). The full pathway, including cross-talk between AHR and other signalling pathways that lead to noncanonical mechanisms of action of AHR and its ligands, overlaid with expression data from experiments 1 (S1 Fig) and 2 (S2 Fig), highlight a total of 28 concordant genes that all achieve differential gene expression at *P*_adj_<0.05. These demonstrate the interplay between the top IPA-identified canonical pathways, with AHR function influencing cell proliferation and estrogen receptor signalling pathways, while heme derivatives biliverdin and bilirubin are known to act as endogenous ligands for AHR [33,34]. Identification of Mitotic Roles of Polo-Like Kinases as a top canonical pathway is indicative of cell proliferative activity that is consistent with *CDC20* and *CDK1* (Table 2) as major hub genes in the network (Fig 3), and with identification of the cyclin-dependent kinase inhibitor *CDKN1A* as the top inhibited upstream regulator (Activation Z-score = −2.764; *P*=5.4×10^-26^) in IPA.

**Figure 4.**
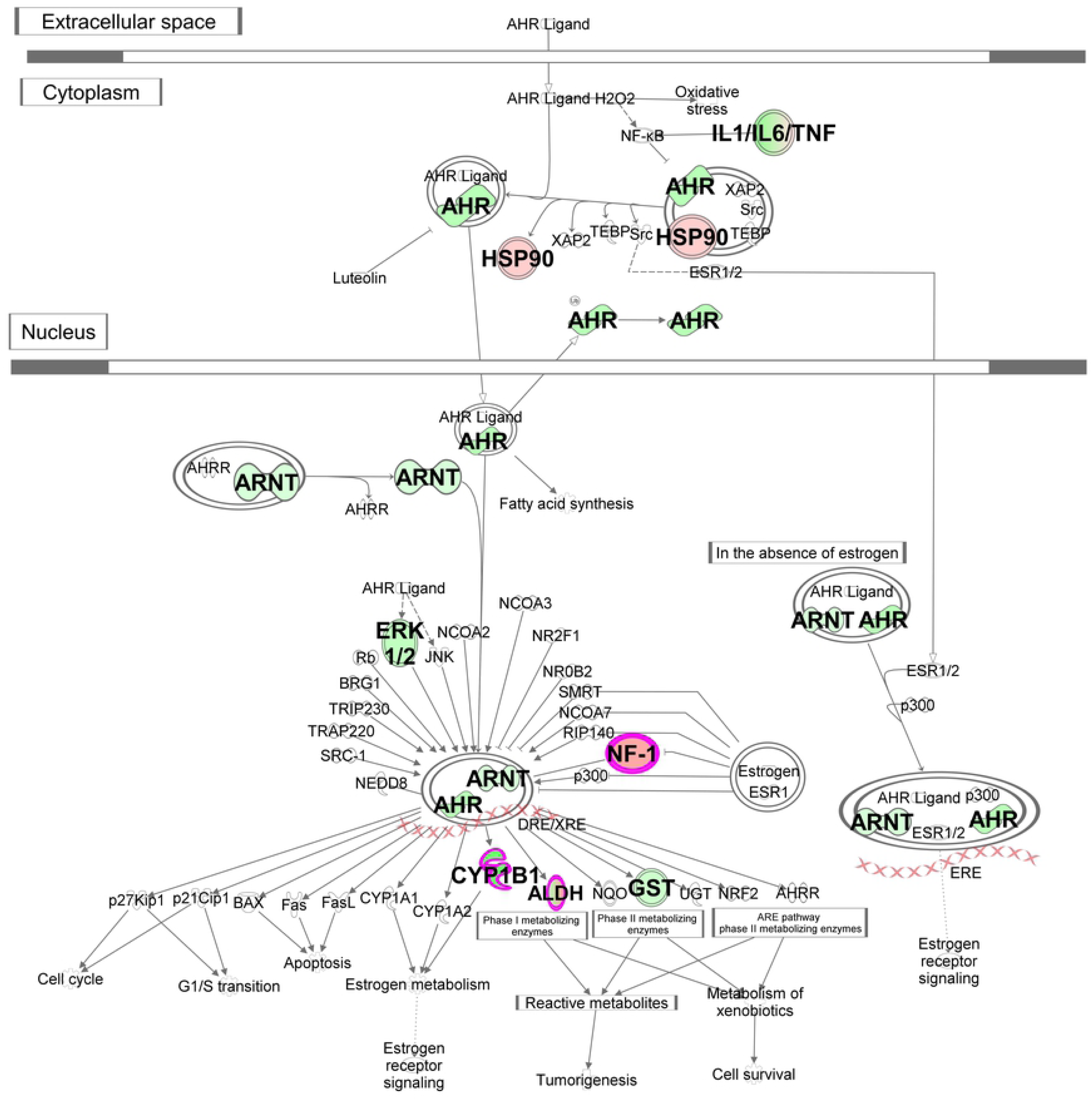
Schematic representation of the core Aryl Hydrocarbon Receptor (AHR) Signalling pathway. The pathway was generated in IPA using data for concordant differentially expressed (*P*_adj_<0.05) genes across the two experiments. Molecules outlined in purple achieved fold-change >2. Genes in green have decreased expression in active cases compared to healthy controls, genes in red have increased expression. The more intense the colour the larger the fold change values. Expression values are based in experiment 1, representative of similar results obtained for concordant genes across the two experiments.

Using Enrichr (S2 Table), signalling pathways involved in cell cycle predominated amongst the top pathways using the Reactome 2016 (“Cell cycle_*Homo sapiens*”, “Cell Cycle, Mitotic_*Homo sapiens*”, and multiple other pathways involved in cell cycle), WikiPathways 2016 (“Cell Cycle *Homo sapiens*”), KEGG 2016 (“Cell cycle_*Homo sapiens*”), and NCI-Nature 2016 (“Aurora B signalling”, “Aurora A signalling” and the “FOXM1 transcription factor network for *Homo sapiens*”, all of which play key roles in cell cycle progression) databases. *CDK1* was also identified as the top PPI Hub Protein using Enrichr (S2 Table). Consistent with our top 10 “induced” gene list, other database comparisons using Enrichr (S2 Table) identified gene sets associated with erythrocyte function including “erythroid cell” (Jensen Tissues Table), “abnormal erythrocyte morphology” and multiple other erythrocyte-related phenotypes (MGI Mammalian Phenotype 2017), “CD71+Early Erythroid” (Human Gene Atlas), “congenital haemolytic anaemia” (Jensen Diseases), and “Haemoglobin’s Chaperone pathway” (BIOCARTA_2016).

### Network, pathway and gene set enrichment analyses comparing active cases and treated cases

Heatmaps were generated for individual expression levels for the top 10 concordant genes expressed at a higher level (Fig 5A), and the top 10 concordant genes expressed at a lower level (Fig 5B), in active cases compared to treated cases in experiment 1. Heatmaps were also generated for the same “induced” and “repressed” probes/genes in experiment 2 (Fig 5C and 5D). In this case, 6/10 and 7/10 top genes from experiment 1 were also in the top 10 most highly differentially expressed genes for “induced” and “repressed” gene sets in experiment 2, respectively, and all achieved fold-change >2 in both experiments. Amongst the 10 most “repressed” genes in experiments 1 and 2 were 3 genes also observed in the comparison of active cases with healthy controls: peptidase inhibitor 3 (*PI3*), as noted above known as an antimicrobial peptide for bacteria and fungi; *ALPL* which encodes an alkaline phosphatase that plays a role in bone mineralization; and *CACNA2D3* which encodes the alpha2delta3 subunit of the voltage-dependent calcium channel complex. Of additional interest in this comparison were “repressed” genes: *CHI3L1* which encodes a chitinase-like protein that lacks chitinase activity but is secreted by activated macrophages and neutrophils; *EMR3* (*ADGRE3*) encoding an adhesion G protein-coupled receptor expressed predominantly in cells of the immune system and playing a role in myeloid-myeloid interactions during inflammation; and *MMP25* that encodes matrix metallopeptidase 25 which inactivates alpha-1 proteinase inhibitor produced by activated neutrophils during inflammation thereby facilitating transendothelial migration of neutrophils to inflammatory sites. Of interest amongst the top 10 “induced” genes in both experiments were: *CXCL10* encoding a chemokine of the CXC subfamily that is a ligand for CXCR3, binding to which results in stimulation of monocyte, natural killer and T-cell migration; *IFNG* encoding interferon-γ, well known for its role in macrophage activation for anti-leishmanial activity; and *GBP1* that encodes a guanylate binding protein induced by interferon.

**Figure 5.**
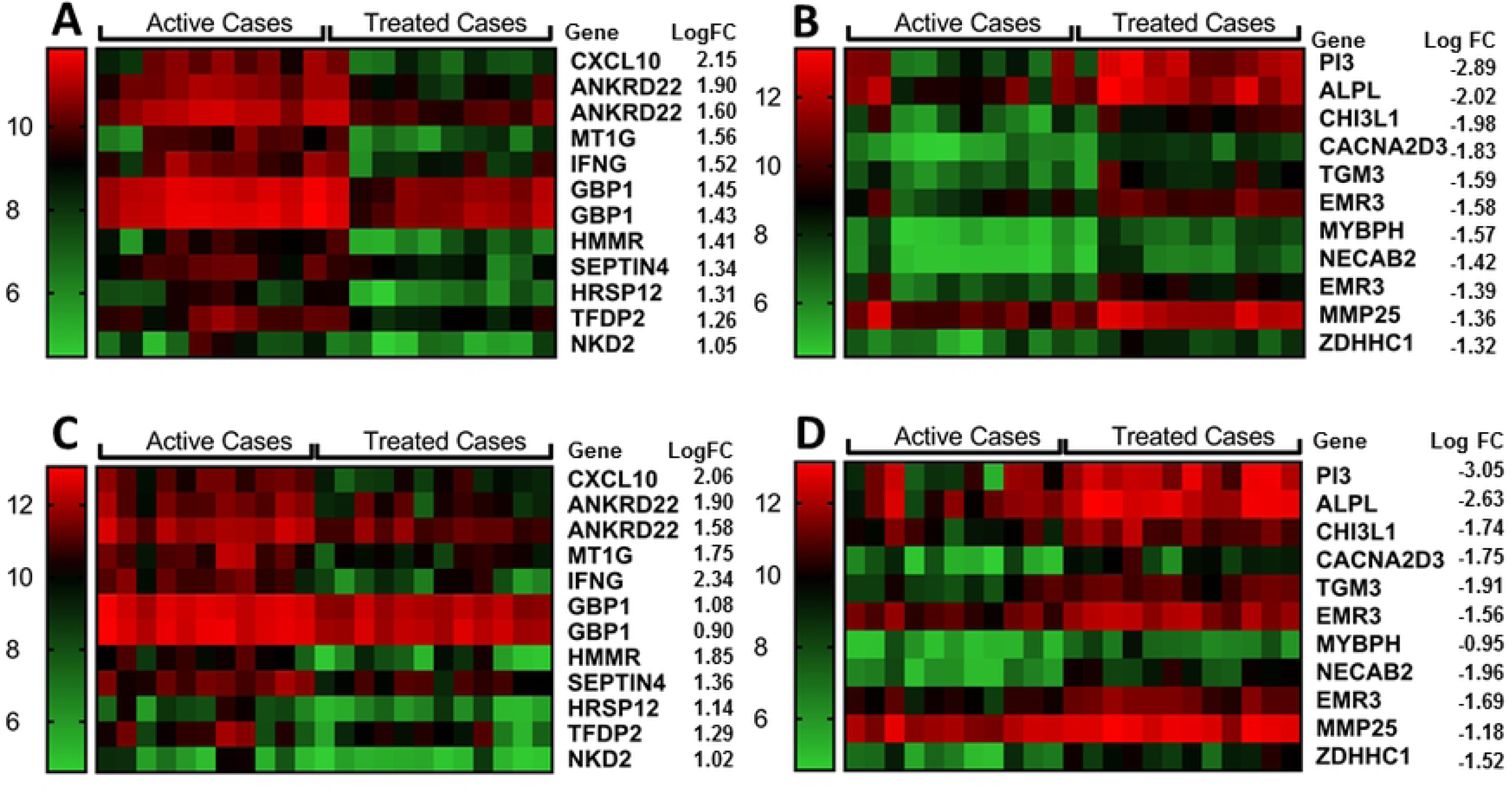
Heatmaps for top differentially expressed genes between active cases and treated cases. (A) top 10 “induced” and (B) top 10 “repressed” genes for differential expression between active cases (N=10) and treated cases (N=10) in experiment 1. (C) and (D) show heatmaps for the same genes in active cases (N=11) and treated cases (N=12) using data from experiment 2. Columns represent individuals and rows represent individual genes, coloured to indicate expression levels based on post-QC normalised and log_2_-transformed data as indicated by the legend to the left of each figure. LogFC = log_2_ fold-change.

As there were only 42 concordant genes that achieved ≥2-fold change in expression, a more global picture of the impact of differential gene expression was obtained by performing IPA and Enrichr analyses using the full set of 210 genes represented by 221 probes that were concordant for differential gene expression at *P*_adj_ ≤0.05. IPA network analysis indicated that 85 of these genes are joined in a single network (Fig 6), with *IFNG* as the major hub gene (i.e. with most connections to other genes in the network), and other major hub genes including *STAT1, SPI1, RARA, NOTCH1* and *MAPK3*. The top canonical pathways included pathogenesis of multiple sclerosis (Table 2), consistent with interconnections between *CXCL10*/*CXCL9*/*CXCL11* and major hub genes *IFNG* and *STAT1* (Fig 6), and the Notch signalling pathway. Aryl hydrocarbon receptor signalling was also identified as a canonical pathway in this analysis at a nominal *P*=0.003 (Table 2). Enrichment for chemokine signalling and Notch signalling pathways were also supported by analyses undertaken using Enrichr (Reactome 2016; WikiPathways 2016, and KEGG 2016 pathways; S3 Table). Consistent with this were top LINCS_L1000_Ligand_Perturbations_Up (S3 Table) for which perturbations of TNFA, IFNG, IL1, IFNA, and HGF were all significant at *P*_adj_<0.01. These ligand perturbations were all associated with differential expression at *CXCL10*, and commonly also at *CXCL11, CXCL9*, and *STAT1*. The major cell types associated with the treatment response were CD14+ monocytes and CD33+ myeloid cell populations (Human Gene Atlas; S3 Table).

**Figure 6.**
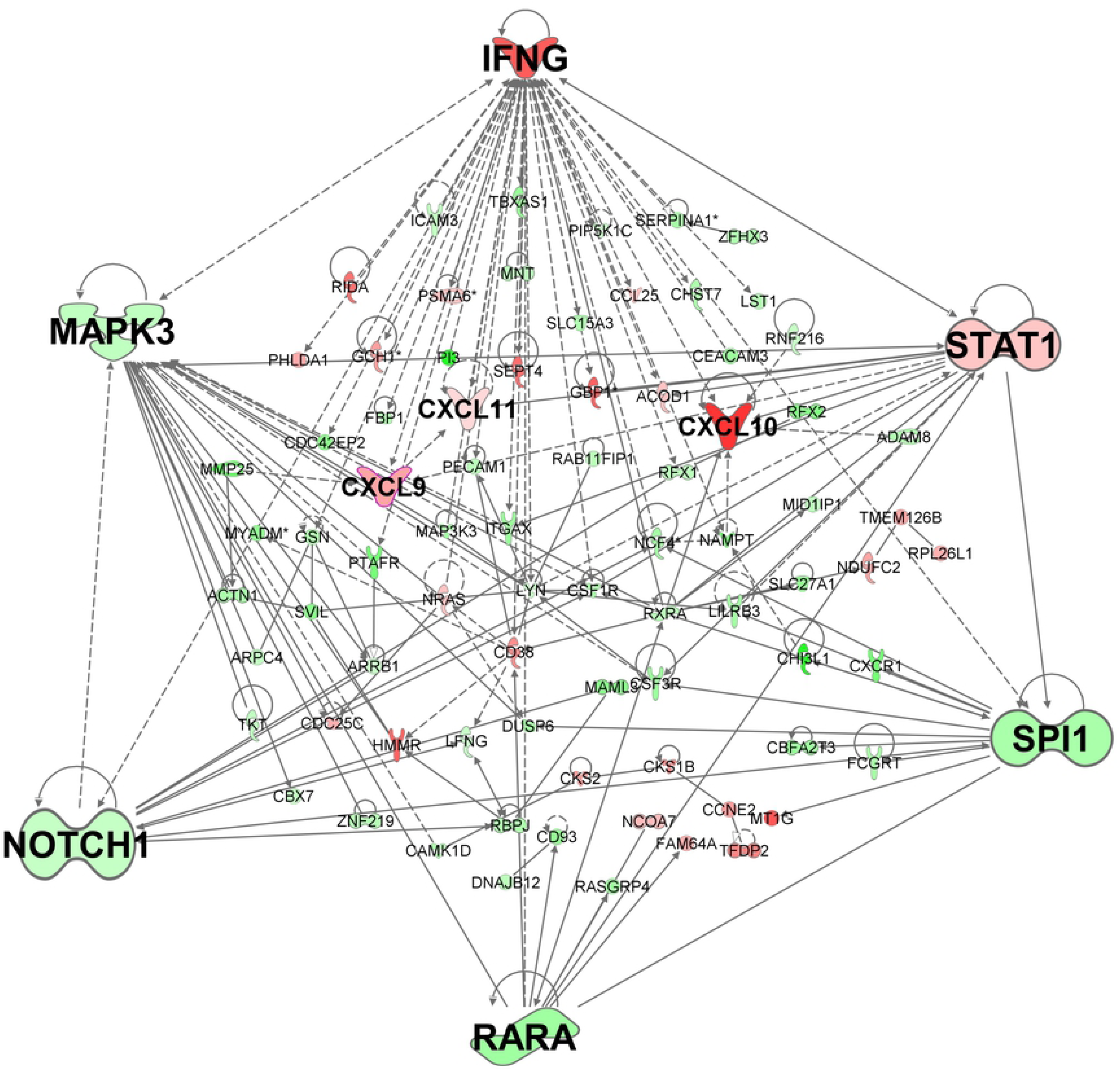
Gene network for concordant genes comparing active cases and treated cases. The network was generated in IPA for 85 (of 210) genes concordant across experiments 1 and 2 for differential expression (adjusted p-value ≤0.05) when comparing active cases and treated cases. Genes in red have increased expression and genes in green have decreased expression when comparing active cases with treated cases. The more intense the colour the larger the fold change values. Expression values are based in experiment 1, representative of similar results obtained for concordant genes across the two experiments.

### Analysis of discordant genes for active cases compared to treated cases

As noted above, we found more differentially expressed probes between treated cases and controls, along with fewer differentially expressed probes between active and treated cases, in experiment 1 compared to experiment 2 (Table 1). We hypothesize that this is due to more effective treatment using liposome encapsulated amphotericin B in experiment 2 compared to the non-liposomal form of the drug employed during experiment 1. We therefore examined the genes that were discordant between active cases and treated cases across the two experiments to understand differences in the cure response. In support of the more efficient cure rate in experiment 2, 7/10 of the top “repressed” genes (namely: *OLIG1, OLIG2, PTGDR2* alias *GPR44, CCR3, CCL23, ALOX15, SLC29A1*) identified as differentially expressed between active cases and treated cases in experiment 2 but not experiment 1 were the same genes that were most repressed in the concordant genes comparing active cases with healthy controls. In comparison, 0/10 of the top “repressed” genes identified as differentially expressed between active cases and treated cases in experiment 1 (but not experiment 2) matched the comparison of concordant genes for active cases and healthy controls. That is, treated cases in experiment 2 were behaving more like healthy controls than were treated cases in experiment 1.

To gain a more global picture of differential gene expression that might inform mechanistic differences in cure rates between the two therapeutic regimes, the 417 genes (from 443 probes) that were differentially expressed between active cases and treated cases in experiment 1 but not experiment 2, and the 988 genes (from 1096 probes) that were differentially expressed between active cases and treated cases in experiment 2 but not experiment 1, were analysed in Enrichr for gene-set enrichment. S4 Table and S5 Table present details of the pathways and gene sets that contrast molecular events that characterise the two different treatment groups. These are summarised in Table 3. For the 988 genes that were differentially expressed between active cases and treated cases in experiment 2 but not in experiment 1 (S4 Table) signalling pathways involved in cell cycle predominated amongst the top pathways using the Reactome 2016 (“Cell cycle_*Homo sapiens*”), Wiki Pathways 2016 (“Cell Cycle *Homo sapiens*”), KEGG 2016 (“Cell cycle_*Homo sapiens*”), and NCI-Nature 2016 (“Aurora B signalling”) databases. In every case there were multiple other pathways involved in cell cycle that achieved Z-scores <-1 and *P*_adj_ ≤0.01. This pattern recapitulates the results obtained in the earlier comparison of concordant genes for active cases and healthy controls (S2 Table), with *CDK1* again identified as the top PPI Hub Protein for this gene set (S5 Table). Consistent with an enhanced rate of cure, multiple immune response signalling pathways (Table 3 and S4 Table) were also identified in this gene set, including IL-1, IL-3, IL-4, IL-6, IL-7 and IL-8 signalling pathways (Reactome 2016, Wiki Pathways 2016, and NCI Nature 2016 databases), Delta-Notch signalling (Wiki Pathways database), chemokine signalling (Wiki Pathways 2016 and KEGG 2016 databases) including specifically IL-8/CXCR2-mediated and CXCR4-mediated signalling (NCI Nature 2016 database), and Fc gamma R-mediated phagocytosis *Homo sapiens* (KEGG 2016 database). Of note, IL-4 was identified as the most significantly down-regulated perturbed ligand pathway (LINCS_L1000_Perturbed_Down; Z-score −1.8, *P*_adj_ 6.36×10^-11^) in this set of genes differentially expressed in active versus treated cases in experiment 2 but not experiment 1. None of these databases showed significant gene set enrichment when interrogated with the 417 genes identified as differentially expressed between active cases and treated controls in experiment 1 but not in experiment 2, i.e. they are not present in Table 3 or S5 Table which compare other enriched gene sets showing differences of interest between experiments 1 and 2. For example, all PPI Hub Proteins identified as significant for the 988 genes that were differentially expressed between active cases and treated cases in experiment 2 but not experiment 1 were related to cell cycle (S5 Table). In contrast, the 5 significant matches to gene sets for PPI Hub Proteins for the 417 genes differentially expressed between active cases and treated cases in experiment 1 but not experiment 2 included the inhibitor of NFκB NFKBIA and the SMAD-signalling pathway gene SMAD9 which transduces signals from members of the TGFβ family. Mutations in NFKBIA are associated with T-cell immunodeficiency [35]. SMAD9 (aliases SMAD8, SMAD8A, SMAD8B, SMAD8/9) transduces signals following ligation of TGFβ family members known as bone morphogenesis proteins (BMPs) to specific BMP (TGFβ family) receptors. Enrichr identified enrichment for a gene set matching genes differentially expressed in BMP4-treated cells (SILAC-Phosphoproteomic Database; *P*=5.4×10^-5^, *P*_adj_=0.003, Z-score = −1.74) from the 417 but not the 988 genes (S5 Table). Another difference was enrichment of the “CD71+Early Erythroid” (Human Gene Atlas) gene set in the 417 genes, while the 988 genes were enriched for gene sets (S5 Table Human Gene Atlas database) associated with B lymphoblasts, CD105^+^ endothelial, CD33+ myeloid, and CD14+ monocytes but not erythroid cells. Overall these analyses of discordant gene sets between experiments 1 and 2 are consistent with our hypothesis that patients treated with a single dose of liposomal amphotericin B (experiment 2) were at a more advanced stage of cure at day 30 post treatment than patients treated with multi-dose non-liposomal amphotericin B (experiment 1).

**Table 3.**
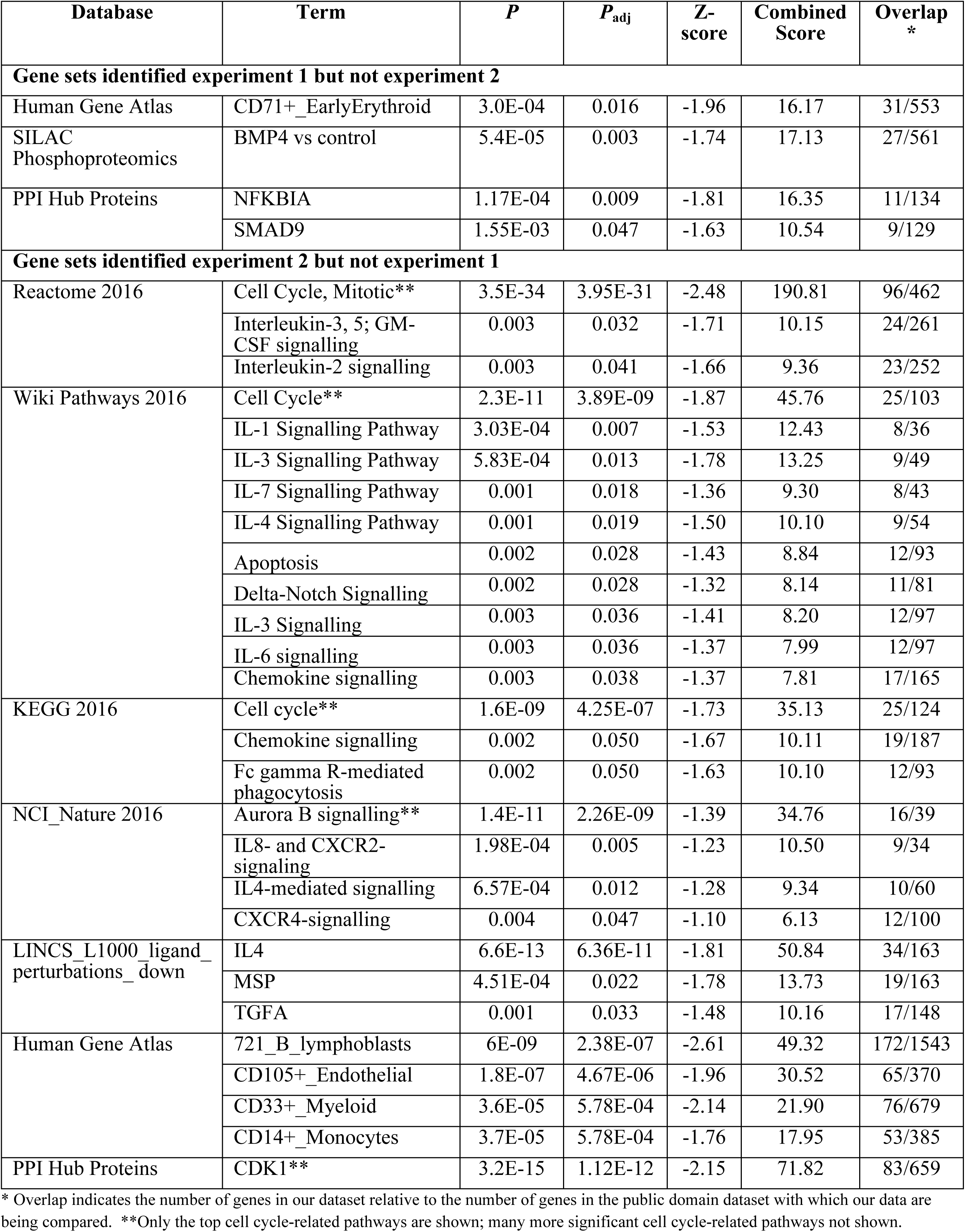
Comparison of Enrichr results for discordant gene sets. The table compares genes sets that were enriched in 417 genes differentially expressed (*P*_adj_ ≤0.05) between active cases and treated cases in experiment 1 but not experiment 2 with those enriched in 988 genes differentially expressed (*P*_adj_ ≤0.05) in experiment 2 but not in experiment 1. Only results where the Z score is <-1 or >1, and the enrichment *P*_adj_ ≤0.05, are included. See methods for an explanation of the combined score. Full data provided in S4 and S5 Tables.

## Discussion

In this study we have analysed whole blood transcriptomic data to further understand the pathogenesis of VL. One original goal of the study was to identify transcriptomic signatures that might differentiate asymptomatic infections from uninfected controls. In our attempt to achieve this we compared both modified quantiferon positive asymptomatic individuals and high antibody positive asymptomatic individuals with healthy endemic controls who were negative for these assays. In the event, we did not find signatures that would be diagnostic for either of these asymptomatic groups compared to negative controls. This was despite longitudinal epidemiological evidence from our study area showing that high antibody individuals are the group at most risk of progressing to clinical VL [6]. Our attention therefore focussed on understanding disease pathogenesis by comparing whole blood transcriptomes from active cases with all healthy controls, and in examining differences in the transcriptome following different regimens of drug treatment. In these comparisons 6 major themes emerged: (i) expression of genes and enrichment of gene sets associated with erythrocyte function in active cases; (ii) strong evidence for enrichment of gene sets involved in cell cycle in comparing active cases with healthy controls (or with more effective cure in experiment 2); (iii) identification of *IFNG* encoding interferon-γ as the major hub gene in concordant gene expression patterns across experiments comparing active cases with healthy controls or with treated cases; (iv) enrichment for interleukin signalling (IL-1/3/4/6/7/8) and a prominent role for CXCL10/9/11 and chemokine signalling pathways in the comparison of active cases with treated cases; (v) the novel identification of AHR signalling as a significant IPA canonical pathway identified from concordant gene expression patterns across experiments comparing active cases with healthy controls or with treated cases; and (vi) global expression profiling support for more effective cure at day 30 post-treatment with a single dose of liposomal encapsulated amphotericin B compared to multi-dose treatment over 30 days.

Interesting in our analysis of top differentially expressed genes and enriched gene sets/pathways between active cases and healthy controls was the predominance of gene sets associated with erythroid cells and function. A recent systematic review [36] found that anaemia has an overall prevalence higher than 90% in VL. Pathogenesis of anaemia based on clinical observations included the presence of anti-erythrocyte antibodies, dysfunction in erythropoiesis, and hemophagocytosis in spleen or bone marrow. Of these, the authors of this review conclude that hemophagocytosis is the most likely cause [36]. The results of our study indicate differential regulation of gene sets associated with abnormal erythrocyte morphology, erythropoiesis, erythrocyte physiology, erythrocyte osmotic lysis, along with decreased haematocrit, spherocytosis and reticulocytosis. The gene sets defining these erythrocyte phenotypes therefore suggest mechanisms other than just hemophagocytosis and could provide important signatures to monitor clinical cure. This is especially relevant given our observation that erythroid related genes were present amongst the discordant genes that were differentially expressed between active cases and cases treated with multi-dose amphotericin B (experiment 1) in which the degree of clinical cure was not as progressed for the same period of treatment with a single dose of liposomal amphotericin B (experiment 2).

A common feature of both the comparison of active cases with healthy controls, and of active cases with treated cases, was the identification of *IFNG* encoding interferon-γ as the major hub gene. This was not itself surprising since interferon-γ plays a key role in activating macrophages to kill *L. donovani* parasites [37]. Studies across the leishmaniases have generally supported the notion that type 1 immune responses and the production of interferon-γ are vital for macrophage activation and parasite elimination [38-40]. It was interesting in our study that transcript levels for *IFNG* were higher in active cases than treated cases, where enhanced interferon-γ responses might have been expected to accompany drug cure. Nonetheless, it concurs with our observations that CD4^+^ T cells in whole blood from active VL patients and treated patients secrete high levels of interferon-γ following stimulation with crude *Leishmania* antigen [41,42], the difference being that only active VL cases secreted IL-10 concurrently with interferon-γ [42]. The higher transcript abundance for *IFNG* in active compared to treated cases in our study suggests return to baseline with treatment in the latter.

Accompanying the central role of *IFNG* as a hub gene when comparing active cases with treated cases was evidence for perturbation of multiple cytokines, including IFNG, IFNA, IL-1, IL-6, and TNF, all of which were supported by differentially expressed gene signatures that generally included CXCL10/11/9 and STAT1. This CXCL10/11/9 chemokine gene expression signature also accounted for the identification of “pathogenesis of multiple sclerosis” [43] as the top disease-related canonical pathway identified using IPA, consistent with a proinflammatory response contributing to disease pathology in active VL. “Pathogenesis of multiple sclerosis” was also identified as a top canonical pathway in spleen tissue and splenic macrophages from *L. donovani* infected hamsters [44], a study in which the authors also noted high interferon-γ expression that was ineffective in directing macrophage activation and parasite killing. STAT1 is a transcription factor activated by ligation of interferon-γ receptors. CXCL10/11/9 are all induced by interferon-γ, all bind to CXCR3, and between them have multiple roles as chemoattractants for monocytes and macrophages, T cells, NK cells, and dendritic cells, and in promoting T cell adhesion. CXCL10 and CXCL9 were also identified as the most highly “induced” genes in comparing lesion transcript profiles with normal skin of patients with American cutaneous leishmaniasis, consistent with their roles in inflammatory cell recruitment [45]. Cxcl9, Gbp1 (encoding the interferon-γ-induced guanylate binding protein *GBP1* identified here as one of the top 10 induced genes when comparing active *versus* treated cases), and Ifng were also identified as part of a common signature of 26 genes upregulated in blood, spleen and liver throughout the course of experimental infection with *L. donovani* in susceptible BALB/c mice, with Cxcl9 and Gbp1 reported as hub genes from a STRING analysis [46].

Given the many studies that have identified the importance of regulatory IL-10 in VL pathogenesis [42, 47-50], it was of some interest in our study that *IL10* was not identified as a top differentially expressed gene or as a significantly enriched signalling pathway in either comparison of active cases with healthy controls, or of active cases with treated cases. Nor did we observed perturbation of IL10R as has been reported in experimental transcriptional profiling studies of VL [46]. Indeed, downregulated expression of the type 2 cytokine gene *IL4* was the strongest response associated with effective cure in liposome-encapsulated amphotericin B treated cases, in line with previous studies showing that IL-4 levels were two-fold higher in VL patients who had failed treatment compared to previously untreated patients, whereas IL-10 levels were comparable in both [49].

One novel observation of our study was identification of AHR signalling as the top canonical pathway when comparing transcriptomes between active cases and healthy controls or treated cases. Through crosstalk between signalling pathways, AHR ligands have been shown to significantly induce IL-10 secretion and inhibit IL-1β and IL-6 production in dendritic cells, and to promote IL-10 production and suppress IL-17 expression in CD4^+^ T cells [30-32]. IL-17 is a potent activator of neutrophils, both through lineage expansion and through their recruitment by regulating chemokine expression. While IL-17 perturbation was not identified in our whole blood transcriptional profiles associated with human VL, evidence from murine models [51] demonstrate a strong role for IL-17 and neutrophils in parasite clearance from liver and spleen. Duthie and coworkers [50] have shown that both IL-10 and IL-17 cytokines are elevated in the serum of active VL patients, reverting to baseline levels with standard antimonial treatments. AHR activation has also been shown to inhibit inflammation through upregulation of IL-22 [52], another cytokine that has been shown to be significantly higher in *Leishmania* antigen stimulated peripheral blood mononuclear cells from active VL cases compared to treated cases [53]. AHR activation during VL may underpin the complex regulation of pro- and anti-inflammatory responses during disease pathogenesis and during response to therapy.

Of potential translational importance in our study was the additional identification of AHR signalling pathway at the top of the Ingenuity “Top Tox List” indicative of its role as a toxic pathology endpoint that could be amenable to therapeutic intervention. AHR locates to the cytoplasm in a stable complex that includes HSP90 observed as a differentially regulated gene in our comparison of active cases with healthy controls. Ligand binding occurs in the cytoplasm and triggers AHR translocation to the nucleus where it binds with ARNT to act as a transcription factor. Both AHR and ARNT were differentially expressed between active VL cases and controls in our study. The AHR response was first associated with xenobiotic induction of metabolizing enzymes, such as the induction of cytochrome P450, family 1, subfamily A, polypeptide 1 (Cyp1a1) following exposure to the polychlorinated dibenzo-p-dioxin 2,3,7,8-Tetrachlorodibenzo-p-dioxin [54]. Multiple AHR ligands are known to induce a “gene battery” of metabolizing enzymes involved in oxidative stress response, cell cycle and apoptosis [55], amongst which are CYP1B1, ALD3B1, ALD3B and ALDH5A1 that were differentially expressed between active VL cases and healthy controls. Transcriptomic profiling of *M. tuberculosis* infected macrophages uncovered evidence for the generation of endogenous AHR ligands through induction of enzymes controlling tryptophan catabolism [56]. The generation of endogenous AHR ligands may likewise explain the role of AHR signalling in VL. For example, heme derivatives biliverdin and bilirubin have both been shown to act as endogenous ligands for AHR, as have arachidonic acid metabolites such as prostaglandins and leukotrienes [33,34]. The former would be consistent with the strong perturbation of erythrocyte function between active VL cases and controls observed in our study. Importantly, addition of exogenous AHR ligands enhanced *M. tuberculosis* infection associated AHR transactivation to stimulate expression of AHR target genes, including IL-1β and IL-23 which stimulate T cell subsets to produce IL-22. This suggests that administration of exogenous ligands could be used as a therapeutic intervention, especially in the knowledge that different exogenous AHR ligands can modulate either regulatory T cell or inflammatory T helper 17 cell differentiation in a ligand-specific fashion to suppress or exacerbate autoimmune disease [57].

Overall, our study has made some novel observations in relation to gene signatures that accompany both active VL disease and clinical cure in treated cases that could provide translatable targets for the development of novel or drug repurposed therapeutic interventions. Furthermore, by studying in more detail the discordant gene patterns that accompanied treatment with single dose liposome encapsulated amphotericin B versus multi-dose non-liposomal amphotericin B we were able to define gene signatures that could be used to monitor progress towards clinical cure.

## Acknowledgements

We would like to thank the hospital staffs at Kala–azar Medical Research Centre, Muzaffarpur for their assistance in the collection of samples and all research scholars of Infectious Disease Research Laboratory, Banaras Hindu University for their kind help during the study.

## Author Contributions

### Conceptualization

Michaela Fakiola, Om-Prakash Singh, Jenefer M. Blackwell

### Data curation

Michaela Fakiola, Om-Prakash Singh,

### Formal analysis

Michaela Fakiola, Om-Prakash Singh, Genevieve Syn, Jenefer M. Blackwell

### Funding acquisition

Shyam Sundar, Jenefer M. Blackwell

### Investigation

Michaela Fakiola, Om-Prakash Singh, Genevieve Syn, Toolika Singh, Bhawana Singh, Jaya Chakravarty

### Project administration

Michaela Fakiola, Om-Prakash Singh, Jaya Chakravarty

### Supervision

Shyam Sundar, Jenefer M. Blackwell

### Writing – original draft

Jenefer M. Blackwell

### Writing – reviewing and editing

Jenefer M. Blackwell, Michaela Fakiola, Om-Prakash Singh.

### Approval of manuscript

All authors read and approved the manuscript.

## Supporting Information

**S1 Table. Demographic and clinical information on study participants**.

**S2 Table. Gene set enrichment analysis for concordant DEGs for active cases versus healthy controls.** The table shows Enrichr results for analysis of 391 genes concordant for differential expression across two experiments comparing active VL cases with healthy controls.

**S3 Table. Gene set enrichment analysis for concordant DEGs for active cases versus treated cases.** The table shows Enrichr results for analysis of 210 genes concordant for differential expression across two experiments comparing active VL cases with treated VL cases.

**S4 Table. Gene set enrichment analysis for discordant genes.** The table shows Enrichr results for analysis of 988 genes differentially expressed between active VL cases and treated VL cases in experiment 2 but not experiment 1.

**S5 Table. Comparison of gene sets enriched for discordant genes.** The table shows Enrichr results for 417 genes differentially expressed between active VL cases and treated VL cases in experiment 1 but not experiment 2, with additional gene sets (see also S4 Table) enriched between active VL cases and treated VL cases in experiment 2 but not experiment 1.

**S1 Fig. Schematic representation of the Aryl Hydrocarbon Receptor (AHR) Signalling in Experiment 1.** The diagram includes all AHR-interacting pathways generated in IPA using data for concordant differentially expressed (*P*_adj_<0.05) genes across the two experiments. Molecules outlined in purple achieved fold-change >2. Genes in green have decreased expression in active cases compared to healthy controls, genes in red have increased expression. The more intense the colour the larger the fold change values. Expression values are based in experiment 1, representative of similar results obtained for concordant genes across the two experiments (see S2 Fig).

**S2 Fig. Schematic representation of the Aryl Hydrocarbon Receptor (AHR) Signalling in Experiment 2.** The diagram includes all AHR-interacting pathways generated in IPA using data for concordant differentially expressed (*P*_adj_<0.05) genes across the two experiments. Molecules outlined in purple achieved fold-change >2. Genes in green have decreased expression in active cases compared to healthy controls, genes in red have increased expression. The more intense the colour the larger the fold change values. Expression values are based in experiment 2, representative of similar results obtained for concordant genes across the two experiments (see S1 Fig).

